# Transcriptome of the coralline alga *Calliarthron tuberculosum* (Corallinales, Rhodophyta) reveals convergent evolution of a partial lignin biosynthesis pathway

**DOI:** 10.1101/2022.03.30.486440

**Authors:** Jan Y Xue, Katy Hind, Matthew A. Lemay, Andrea Mcminigal, Emma Jourdain, Cheong Xin Chan, Patrick T. Martone

## Abstract

The discovery of lignins in the coralline red alga *Calliarthron tuberculosum* raised new questions about the deep evolution of lignin biosynthesis. Here we present the transcriptome of *C. tuberculosum* supported with newly generated genomic data to identify gene candidates from the monolignol biosynthetic pathway using a combination of sequence similarity-based methods. We identified candidates in the monolignol biosynthesis pathway for the genes 4CL, CCR, CAD, CCoAOMT, and CSE but did not identify candidates for PAL, CYP450 (F5H, C3H, C4H), HCT, and COMT. In gene tree analysis, we present evidence that these gene candidates evolved independently from their land plant counterparts, suggesting convergent evolution of a complex multistep lignin biosynthetic pathway in this red algal lineage. Additionally, we provide tools to extract metabolic pathways and genes from the newly generated transcriptomic and genomic datasets. Using these methods, we extracted genes related to sucrose metabolism and calcification. Ultimately, this transcriptome will provide a foundation for further genetic and experimental studies of calcifying red algae.

## Introduction

Coralline red algae (Corallinales, Sporolithales, Hapalidiales) are a diverse lineage of calcified seaweeds that play important ecological roles in nearshore ecosystems worldwide: they stabilize coral reefs by creating a calcium carbonate matrix [1–3], induce settlement of invertebrate taxa [4–6], and contribute to the storage of blue carbon through the creation of biogenic calcium carbonates [7, 8]. In recent years, there has been increased global attention paid to coralline algae. Taxonomists are clarifying their vastly underestimated species diversity [9–12]; ecologists and physiologists are documenting interspecific variation in coralline growth and calcification, particularly in response to climate stress, which may ultimately impact marine communities [13–17]; evolutionary biologists are examining patterns in coralline trait evolution [18–20] and using >100 million-year-old coralline fossils to strengthen modern phylogenies [21, 22].

The discovery of lignins within cell walls of the coralline species *Calliarthron cheilosporioides* (Corallinales, Rhodophyta) dramatically changed our perspective on the evolution of lignin biosynthesis [23]. Lignins are complex aromatic polymers predominantly found in the secondary cell walls of plant support tissues [24, 25] and were long considered to have evolved when land plants emerged from the oceans, enabling upright growth in air [26]. Among the principal chemical components of wood, lignins in plant secondary cell walls help reinforce tissue mechanical properties, permit hydraulic transport, and increase pathogen resistance [27, 28]. In the articulated coralline *C. cheilosporioides*, lignins were found predominantly within decalcified flexible joints, called genicula [23], that have remarkable biomechanical properties, permitting this articulated coralline species to thrive along wave-battered coastlines [29, 30].

Because lignin biosynthesis is physiologically complex and involves several enzymes in the monolignol pathway [31–33], Martone et al. [23] proposed that much of the lignin biosynthetic pathway may have predated land plants altogether, evolving in a common ancestor of red and green algae more than one billion years ago. Alternatively, some (or all) of the monolignol biosynthetic pathway may have evolved independently in the embryophyte and rhodophyte lineages. For example, one important enzyme involved in S-lignin production (F5H) evolved independently in lycopods and embryophytes [34, 35]. Moreover, candidate genes related to monolignol biosynthesis have since been found in diverse algal lineages such as diatoms, dinoflagellates, haptophytes, cryptophytes, and green and red algae [36], raising questions about how the monolignol pathway may have evolved across such evolutionarily divergent lineages. Until now, questions about monolignol evolution have largely gone unanswered as transcriptomic and genomic data have mostly been limited to non-coralline red algae (e.g. [37–40] but see [41]).

Here we present a transcriptome of the articulated coralline *Calliarthron tuberculosum* (a sister species of *C. cheilosporioides*) to investigate the evolutionary history of monolignol biosynthesis. Additionally, though a complete mitochondrial genome [42] and a draft nuclear genome [43] of *C. tuberculosum* were previously published, herein we generated a revised nuclear genome assembly using new short-read sequence data to aid validation of transcriptomic reads. Based on comparative analysis of genome and transcriptome data, we identify gene candidates for a putative monolignol biosynthetic pathway in *C. tuberculosum* and investigate evolutionary relationships of these enzymes with those from other taxonomic groups, including their land plant counterparts. We also provide a list of annotated genes in the *C. tuberculosum* transcriptome and a simplified method for extracting genes from metabolic pathways. We illustrate the utility of this dataset by extracting gene candidates involved in sucrose metabolism and calcification. This transcriptomic dataset provides a foundation for future studies of coralline algal ecology, physiology, and evolution.

## Results

### The C. tuberculosum transcriptome is complete and supported by genomic data

Two transcriptomic datasets were generated from *Calliarthron* thalli: one from whole tissue (calcified intergenicula plus uncalcified genicula; sample I+G/PTM1 in the deposited data) and a second from intergenicular (i.e., calcified) tissue only (sample I/PTM2). Transcriptome sequencing based on RNA-Seq produced 38.8 total Gb of sequence data (17.3 Gb for sample I+G; 21.5 Gb for sample I). Reads were assembled *de novo* using Trinity. The whole tissue dataset had 172,700,376 total reads and the intergenicular tissue dataset had 215,491,160 total reads with an overall average coverage of 677-fold. A third reference transcriptome combining data from both tissues was assembled independently. All three datasets were combined for subsequent analysis to increase coverage and maximize discovery. The transcriptome data were considered complete based on the recovery of core eukaryotic genes (e.g. 94.5% of CEGMA and 87.8% of BUSCO genes based on TBLASTN; Fig S1A). Genomic sequences were also assembled for *C. tuberculosum* (Table S1), but these remain highly fragmented and were used only as additional support to the transcriptome data in subsequent searches below. More than half (18840; 56.6%) of the 33301 transcripts in the reference transcriptome were supported by the genome data (BLASTN, *E* ≤ 10^-5^).

### The incomplete monolignol biosynthetic pathway in Calliarthron tuberculosum

The combined *C. tuberculosum* transcriptomic dataset was searched for genes encoding enzymes from the monolignol biosynthetic pathway. The transcriptomic dataset was translated into all six reading frames and queried with a combination of homology-based approaches, including HMMER searches and KEGG based annotations. Closest homologs from *Arabidopsis thaliana* were also verified (BLASTN, *E* ≤ 10^-30^). We identified gene candidates of 4CL, CCR, CAD, CSE, and CCoAOMT, but not HCT, COMT, PAL, TAL, or PTAL (Fig 1). PAL/TAL/PTAL was considered absent as only fragmented (and no full length) sequences were identified. Evidence for the presence of homologous p450 enzymes (C3H, C3H, and F5H) was weak; as a result, their status was classified as ambiguous (Fig 1). All sequences identified had genomic support (BLASTN, *E* ≤ 10^-5^) except for those identified for PAL/TAL/PTAL.

**Fig 1.**
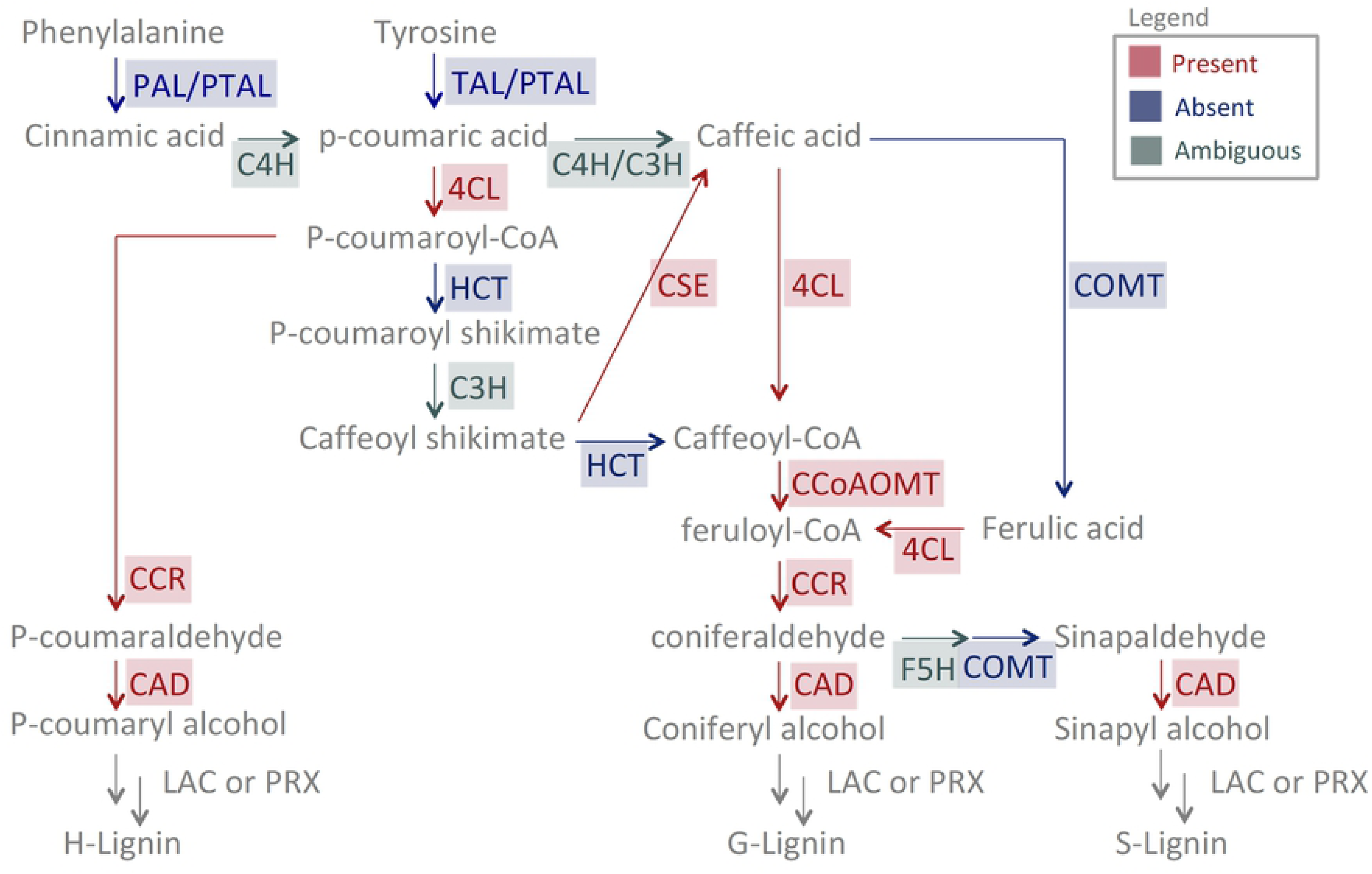
The presence of *C. tuberculosum* sequence candidates in the monolignol pathway. Red indicates presence of a putative homolog in *C. tuberculosum*; blue indicates no significant hits; green indicates ambiguous presence. Note how the PTAL/PAL/TAL sequences obtained from the HMMER search were indicated as absent as all sequences found were too short, 1/4-1/3 in length relative to those in land plants. All sequences identified have genomic support except for PTAL/PAL/TAL.

Candidate sequences from *C. tuberculosum* (bolded as contig_gene_isoform in Figs 2, 3, and 4) were characterized by comparing key residues with their land plant homologs in multiple sequence alignments. The evolutionary relationships between the identified *C. tuberculosum* sequences, closely related sequences in additional taxa, and sequences from the broader protein family of their land plant homologs were analyzed in gene trees. Below we describe in detail results for the main biosynthetic enzymes 4CL, CCR, and CAD (Figs 2, 3, and 4). Descriptions of the other biosynthetic enzymes CCoAMT, CSE, and the cytochrome P450 sequences C3H, C4H, F5H are found in Appendix S1 and Figs S2-S4.

**Fig 2.**
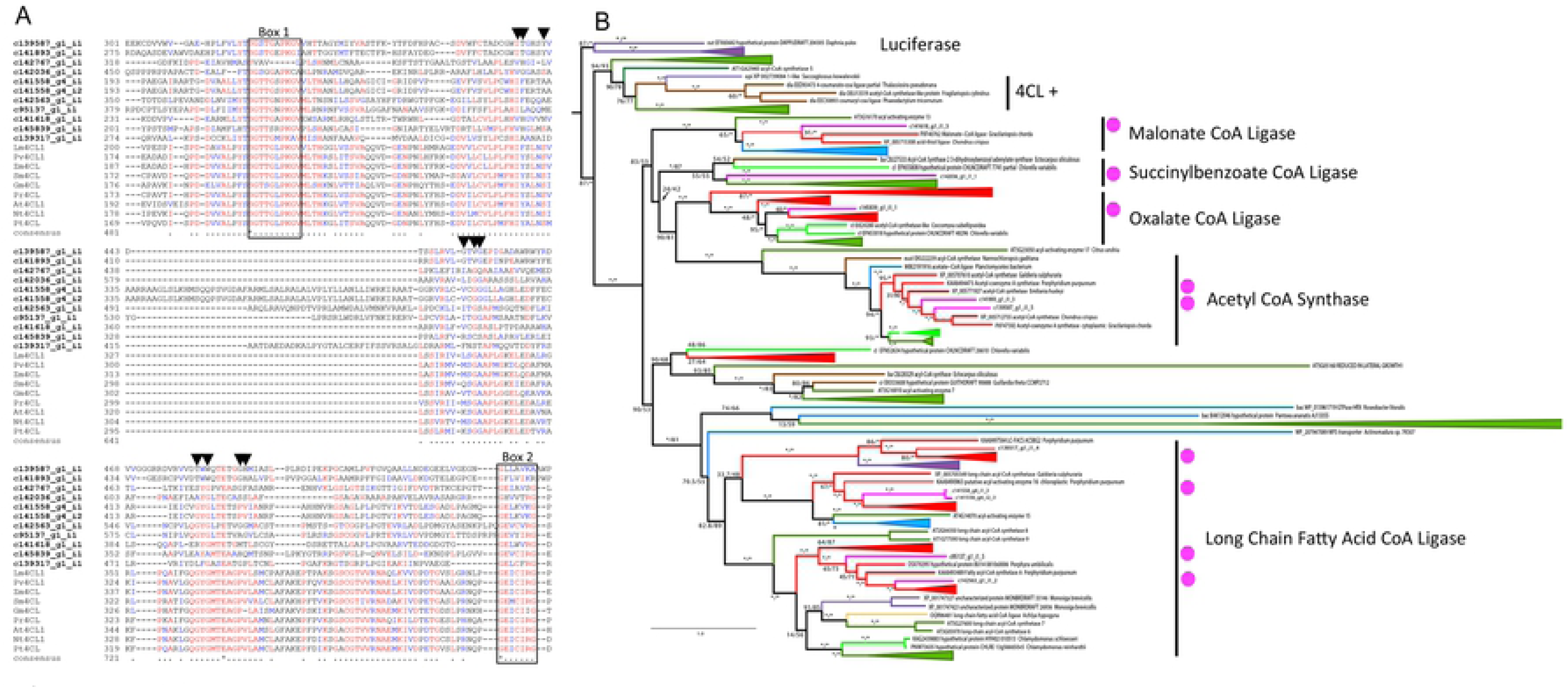
4CL candidates from *C. tuberculosum* in relation to plants and other taxa. **(A)** Partial alignment of *C. tuberculosum* candidates (bolded) and embryophyte 4CL sequences. Residues involved in hydroxycinnamate binding are indicated with black triangles [61, 62]. Phenylalanine substrate binding pocket is indicated with Box I and Box II. **(B)** Maximum likelihood acyl-activating enzyme (AAE) gene tree showing relationships between *Calliarthron* sequences (magenta dots) and other taxa (Embryophyta – dark green, Chlorophyta – light green, Rhodophyta – red, Animalia and Opisthokonta – purple, Bacteria and Cyanobacteria – blue, Oomycota, Mycetozoa and Fungi – yellow, Ochrophyta – brown). Functionally demonstrated plant 4CLs are labelled (+). Additional functional groups are labelled [44, 45]. Ultrafast bootstrap values > 95 are marked by *. Model = WAG+F+G4. Sites with ≤ 80% occupancy were removed. Accession numbers can be found in Appendix S1.

**Fig 3.**
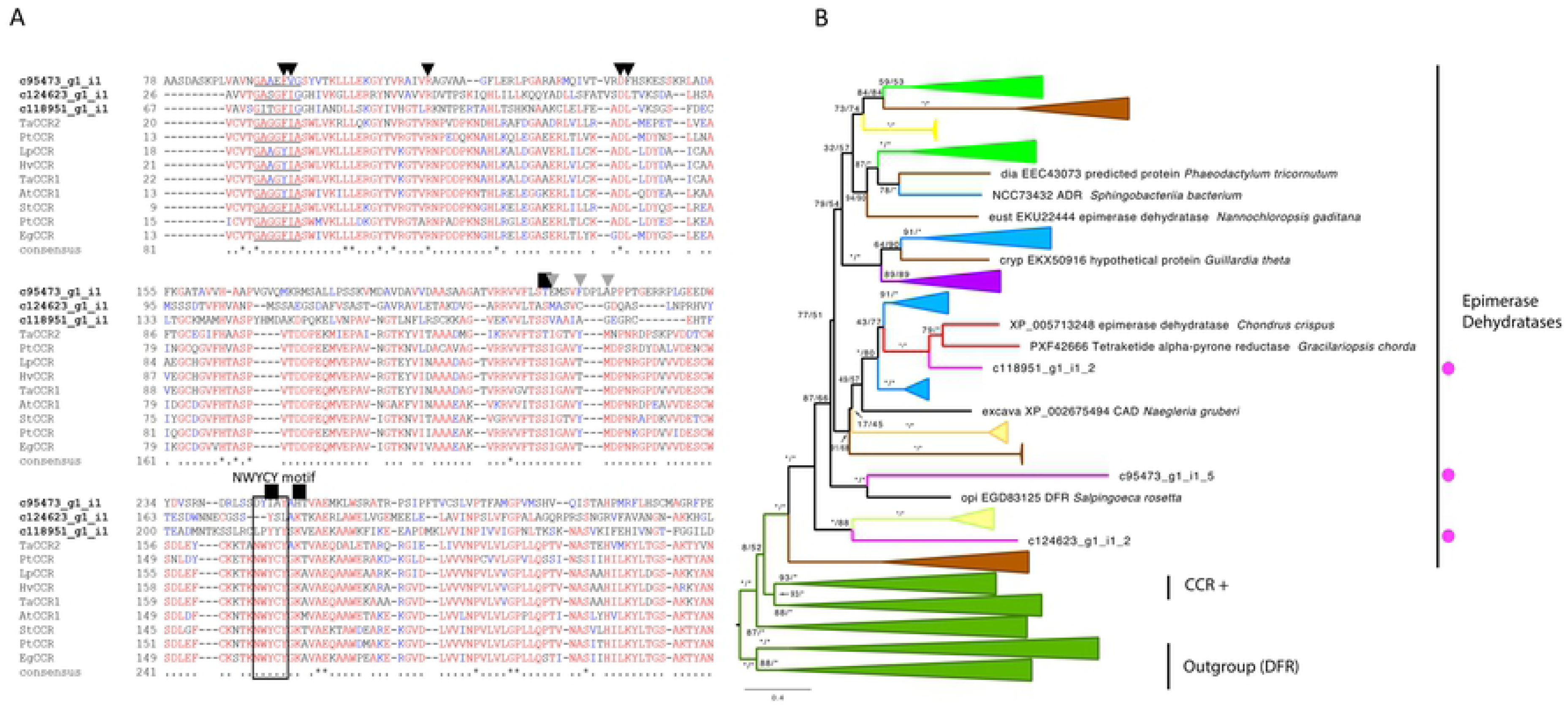
CCR candidates from *C. tuberculosum* in relation to plants and other taxa. **(A)** Partial alignment *C. tuberculosum* candidates (bolded) and land plant CCR sequences. Catalytic residues are labelled with NWYCY [64] and additional residues are indicated above with a black box. NADPH binding pocket residues are indicated with black triangles [65] and the GXXGXX[A/G] motif is underlined [66]. Hydroxycinnamonyl binding pocket residues are indicated with a gray triangle [65]. **(B)** CCR maximum likelihood gene tree showing relationships between *C. tuberculosum* (magenta dots) and other taxa (Embryophyta – dark green, Chlorophyta – light green, Rhodophyta – red, Animalia and Opisthokonta – purple, Bacteria and Cyanobacteria – blue, Oomycota, Mycetozoa and Fungi – yellow, Ochrophyta – brown). Functionally demonstrated plant CCRs are labelled (+). Additional functional groups are labelled. Ultrafast bootstrap values >95 are marked by *. Model = LG+G4. Sites with ≤ 80% occupancy were removed. Accession numbers can be found in Appendix S1.

**Fig 4.**
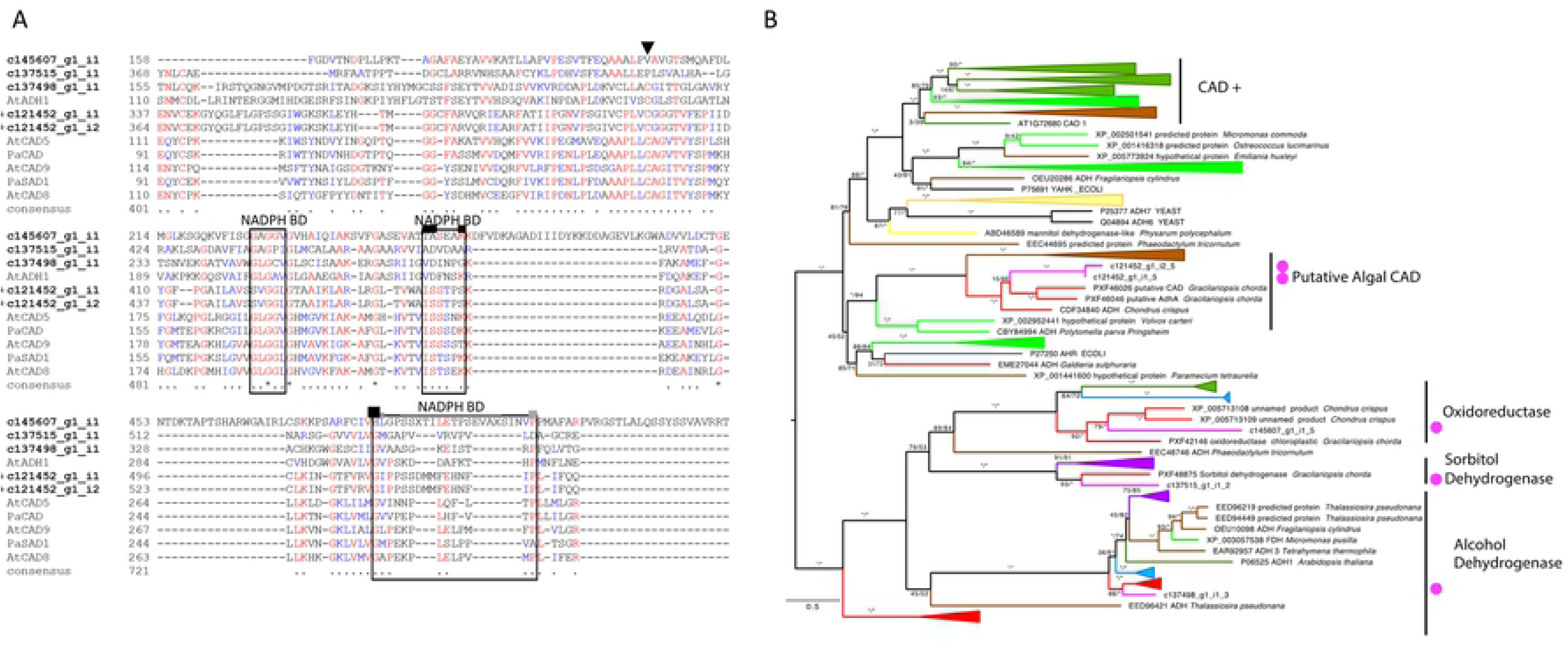
CAD candidates from *C. tuberculosum* in relation to plants and other taxa. **(A)** Partial alignment of *C. tuberculosum* CAD sequence candidates (bolded) with land plant CAD sequences. Zn^+2^ ion coordinating and proton shuttling residues are indicated with the black triangle, NADPH or NADH interacting residues are boxed. Hydrostatic interaction forming residues are indicated with a black box. Putative substrate-binding residues are indicated with grey boxes. [67–69] **(B)** CAD maximum likelihood gene tree showing relationships between *C. tuberculosum* (magenta dots) and other taxa (Embryophyta – dark green, Chlorophyta – light green, Rhodophyta – red, Animalia and Opisthokonta – purple, Bacteria and Cyanobacteria – blue, Oomycota, Mycetozoa and Fungi – yellow, Ochrophyta – brown). Alcohol dehydrogenase (ADH) sequences from yeast, and aldehyde reductase (YAHK and AHR) sequences from *E. coli* were used as the ADH family is closely related to that of CAD [70, 71]. Functionally demonstrated plant CADs are labelled (+). Additional functional groups are labelled. Ultrafast bootstrap values >95 are marked by *. Model = LG+G4. Sites with ≤ 80% occupancy were removed. Accession numbers can be found in Appendix S1.

### Identification of 4CL candidates

4CL is an acyl-CoA synthase in the monolignol pathway and a member of the acyl-activating enzyme (AAE) superfamily. 4CL converts p-coumaric acid, caffeic acid, and ferulic acid into their respective hydroxycinnamoyl-CoA thioesters. We identified 11 candidate 4CL-coding transcripts: two based on KEGG analysis and nine additional sequences based on HMMER searches (Fig 2A). A query of these sequences against the *A. thaliana* proteome returned related proteins within the acyl-activating enzyme superfamily but not the *A. thaliana* 4CL (Table S2). Moderate sequence conservation exists in substrate binding and hydroxycinnamate binding residues between 4CL candidates in *C. tuberculosum* (bolded) and 4CLs in land plants (identity similarity [IS] > 70% Fig 2A).

In the 4CL gene tree analysis, most *C. tuberculosum* sequences grouped with sequences from other Rhodophytes (Fig 2B). In addition, *C. tuberculosum* sequences grouped within several functional clades including malonate CoA ligase (ultrafast bootstrap support [BS] = 100%), succinylbenzoate CoA ligase (BS = 87%), oxylate CoA ligase (BS = 100%), acetyl CoA synthase (BS = 100%), and the long chain fatty acid CoA ligase (BS = 89%) (magenta dots, Fig 2B) [44, 45]. In contrast, embryophyte 4CL sequences form a clade separated from candidate 4CL sequences in *C. tuberculosum* (BS = 99% Fig 2B) by the luciferase containing outgroup. Thus, 4CL candidates in *C. tuberculosum* did not show any clear homology to functionally demonstrated 4CL sequences from embryophytes.

### Identification of CCR candidates

CCR is the first committed enzyme in the monolignol pathway, reducing cinnamoyl-CoA esters to cinnamaldehydes. We identified three sequences as candidate CCR-coding transcripts: one based on KEGG analysis and two additional sequences based on HMMER searches (Fig 3A). A query of these sequences against the *A. thaliana* proteome returned sequences within the CCR family (CCR7, CCR4, CCR-Like6) (Table S2). Substrate-binding residues (NWYCY) and the hydroxycinnamonyl-binding pocket showed low sequence conservation (IS <80%). In contrast, the core catalytic residues (S, T, and K) and NADPH-binding residues appear to be conserved (IS >90%) between the candidate sequences in *C. tuberculosum* and CCRs in land plants (Fig 3A).

In the CCR gene tree analysis, *C. tuberculosum* sequences varied in their relatedness to other taxa with some sequences closer to Rhodophytes and others more closely related to Oomycota/Mycetozoa/Fungi (Fig 3B). Additionally, CCR candidates in *C. tuberculosum* were mapped with epimerase dehydratase type sequences that included the *A. thaliana* CCR family (Fig 3B). Sequences from *C. tuberculosum* grouped with epimerase dehydratase type sequences of non-embryophyte origin. In contrast, embryophyte CCR, class 2 CCR, and CCR-like form an independent clade (BS >97%). The embryophyte CCR clade and the non-embryophyte epimerase dehydratase clade (containing sequences from *C. tuberculosum*) were more closely related than the embryophyte dihydroflavonol-4-reductase protein (DFR) group within the overall epimerase dehydratase family.

### Identification of CAD candidates

CAD, the final step in the monolignol pathway, is an alcohol dehydrogenase converting various hydroxycinnamaldehydes to their respective hydroxycinnamyl alcohols. SAD, proposed to catalyze this same reaction for sinapyl monolignols [46], is added into our analysis despite debate over their function. We identified five sequences as candidate CAD-encoding transcripts: two based on KEGG analysis and three additional sequences based on HMMER searches (Fig 4A). A query of these sequences against the *A. thaliana* proteome returned CAD2 and other alcohol dehydrogenases (Table S2). NADPH-binding motifs show moderate conservation (IS >80%) (Fig 4A). One *C. tuberculosum* sequence showed high conservation with land plant counterparts, suggesting a promising CAD candidate (+ in Figs 3A and 3B).

In the CAD gene tree analysis, all *C. tuberculosum* sequences grouped with sequences from other Rhodophytes (Fig 4B). CAD candidates in *C. tuberculosum* were mapped with their embryophyte CAD counterparts and closely related alcohol dehydrogenases. Sequences from *C. tuberculosum* grouped together with oxidoreductases (BS = 100%), sorbitol dehydrogenases (BS = 100%), general alcohol dehydrogenases (BS = 100%), and an algal CAD clade (BS = 100%). Sequences in this algal CAD clade were based on previous sequence similarity-based annotation and have not been functionally demonstrated. In contrast, the land plant CAD and SAD sequences form their own clades (BS 100%; Fig 4B) that are separated from the *C. tuberculosum* candidates by the functionally distinct alcohol dehydrogenases, such as yeast alcohol dehydrogenase 7 (ADH7) and *E. coli* aldehyde reductase (YAHK).

### Identification of additional metabolic pathways in Calliarthron tuberculosum

To enable broad and rapid identification of *C. tuberculosum* genes involved in specific metabolic processes, we present two general tools for gene identification within the *C. tuberculosum* transcriptome dataset using KEGG based annotations. This involves extracting whole metabolic pathways or individual genes (see Appendix S1; Fig S5). We included annotations for all metabolic genes recovered in the *C. tuberculosum* transcriptome (Table S3). We identified 36 putative *C. tuberculosum* genes present in the starch and sucrose metabolism pathway (Fig S5; Table S4). In addition, we individually searched for genes potentially involved in calcification [41,47,48] and identified 13 sequence candidates related to calcium transport, six related to inorganic carbon transport, five related to pH homeostasis, 19 putative carbonic anhydrases, and 12 putative HSP90 genes (Table S5).

## Discussion

### Evidence for convergent evolution of monolignol biosynthesis

Using sequence similarity methods with genes from the monolignol pathway in land plants, we identified candidates for five genes related to monolignol biosynthesis (4CL, CCR, CAD, CCoAOMT, and CSE) from the newly generated *C. tuberculosum* transcriptomic dataset. These gene candidates are supported by genomic evidence, retain major motifs from their respective gene family, and return their *A. thaliana* counterpart in reciprocal BLAST analyses, suggesting that these enzymes may function similarly in monolignol biosynthesis in *C. tuberculosum*.

Despite supporting evidence from sequence similarity analyses, functional predictions for candidate sequences in the monolignol pathway within *C. tuberculosum* are obscured by the gene tree analysis. If the monolignol pathway in embryophytes and *C. tuberculosum* evolved in a common ancestor and was retained through conserved evolution, we would expect their sequences to form functional clades uninterrupted by functionally divergent protein sequences. However, with the exception of the CCoAOMT candidate, our gene tree analyses consistently showed that monolignol biosynthetic genes in land plants are not sister to those in *C. tuberculosum*. *C. tuberculosum* sequences were found within each respective overall protein family, but consistently grouped with land plant genes of non-monolignol forming function. If these *C. tuberculosum* sequences are functionally homologous to the monolignol biosynthesis counterpart in land plants, then they likely arose independently in *C. tuberculosum*. Convergent evolution in protein function, with phylogenetic patterns of protein sequences with similar functions intersected by sequences with dissimilar functions, is not uncommon in cell wall synthesizing enzymes [49]. Biosynthetic enzymes in *C. tuberculosum* could have evolved similar substrate specificity after the divergence of red algae and land plants or, alternatively, may reflect genes that were individually acquired. Previous evidence suggests that the core monolignol biosynthesis genes (4CL, CCR, and CAD) in *C. tuberculosum* may have been acquired through horizontal gene transfer from a bacterial source [36]. Thus, over evolutionary time genes in *C. tuberculosum* may have developed enough synchronicity in gene expression and protein regulation to produce an ad hoc monolignol biosynthetic pathway.

Alternatively, the phylogenetic evidence might suggest that gene candidates in *C. tuberculosum* do not function in monolignol biosynthesis and instead have a function similar to their sister sequences within their distinct phylogenetic groupings. For example, considering only clustering patterns in the phylogenetic data, perhaps *C. tuberculosum* contig 141618 functions as a CoA ligase that acts on malonate and not coumarate (4CL enzyme) (Fig 2B). However, the tandem use of stricter curated sequences in our predictive HMM models and more flexible HMM models with previously annotated data, such as KEGG annotations, improves our confidence in finding potential gene candidates. Biochemical or functional assays will ultimately be needed to verify the function of candidate gene sequences.

### The monolignol biosynthesis pathway and missing steps in Calliarthron tuberculosum

Several key steps in the monolignol biosynthetic pathway were not recovered in the *C. tuberculosum* transcriptome, including PAL, TAL, PTAL, HCT, COMT, C3H, C4H, or F5H. Although we cannot dismiss that these observations may be due to fragmented sequences in the assembled genome and transcriptome data, we present several other possibilities.

The ammonia-lyase PAL, TAL, or PTAL creates the first substrates in the monolignol biosynthetic pathway [50–52]. Although no full-length homologs were identified in the *C. tuberculosum* transcriptome, short sequence candidates identified may represent a fragmented gene. However, these short sequences lacked genomic support, indicating they may be contaminants of non-*Calliarthron* origin. For this reason, PAL, TAL, and PTAL are currently indicated as absent (Fig 1). If these are indeed from *C. tuberculosum*, RACE amplification could help determine if the short ammonia-lyase we identified has a longer transcript. *C. tuberculosum* likely has an ammonia-lyase acting on phenylalanine or tyrosine since PAL and TAL are also key enzymes in producing flavanoids and coumarins, which have been previously detected in both fleshy and coralline red algae [53]. Further validation will be required to elucidate their presence.

C3H, C4H, or F5H are p450 monooxygenases responsible for converting substrates across the monolignol pathway eventually resulting in H to S to G type monolignols, respectively (Fig 1). P450 sequence candidates have been identified, but their substrate-specific identity as C3H, C4H, or F5H homologs is unclear. The cytochrome P450 sequence candidates from the *C. tuberculosum* transcriptome form two divergent groups. One group is likely involved in carotenoid biosynthesis, positioned within the CYP97 clade, while the other group forms their own clade of unknown function (Fig S2B). The identified candidates from *C. tuberculosum* may have multi-substrate specificities, acting on various substrates, including monolignol intermediate products. Some substrate promiscuity has previously been observed within members of the cytochrome P450 enzyme family [54, 55]. Alternatively, each of the identified P450 clades in *C. tuberculosum* could contain a new class of cytochrome P450 capable of functioning in H-, G-, or S- unit monolignol biosynthesis. This proposed convergent evolution of a distinct and independently-evolved cytochrome P450 involved in monolignol production has previously been documented in the clubmoss *Selaginella moellendorffii* (F5H) [34, 35]. In any case, the presence of unique P450s represents an interesting avenue of exploration to elucidate substrate specificity and functionality in the monolignol pathway in *C. tuberculosum*.

HCT is one alternative route shifting monolignol synthesis from H- to G- to S- types using a temporary shikimate decoration (Fig 1) [56]. Its absence could suggest that *C. tuberculosum* does not utilize an HCT enzyme or create G lignin using this route. Another alternative route in G- and S- type monolignol synthesis utilizes a CSE enzyme that acts on caffeoyl shikimate, an HCT downstream product (Fig 1). The absence of an HCT is at odds with the CSE enzyme identified in this study (Fig 1), suggesting that the CSE candidate identified may not be utilized in the monolignol biosynthetic pathway for *C. tuberculosum*. Though this absence could be due to fragmentation in the transcriptome, more data are required for further validation.

COMT is necessary for S type monolignol production in angiosperms [57–59]. The absence of this enzyme raises questions about how *C. tuberculosum* can produce sinapyl alcohol, a precursor component for S monolignols. Some evidence exists for a bifunctional enzyme in pine that can function as both COMT and CCoAOMT (named AEOMT) in heterologous systems [60]. However, only moderate-to-low sequence similarity is shared among CCoAOMT, COMT, and the bifunctional AEOMT. Perhaps a similar protein with broad substrate specificity is present in *C. tuberculosum* but has yet to be identified based on sequence similarity.

## Conclusion

In summary, we have identified several gene candidates in the *C. tuberculosum* transcriptome that represent central components in the monolignol biosynthetic pathway, helping to explain the surprising presence of lignins in this coralline red alga. Despite the complexity of monolignol biosynthesis, and contrary to the predictions outlined in Martone et al. [23], our gene trees do not demonstrate a deeply conserved evolution of monolignol biosynthesis, but instead suggest that each of the enzymes identified in *C. tuberculosum* likely evolved independently from those found in land plants. Interestingly, there remain several key enzymes in the monolignol pathway whose sequences have not been identified, including those related to pathway entry and to shifting the types of monolignols produced that would form H-, G-, and S-lignins within the cell wall. Further biochemical evidence and validation of sequence expression will be necessary to provide functional support for both the genes identified and to elucidate potential alternative routes in the monolignol biosynthetic pathway in *C. tuberculosum.* By providing methods to easily identify additional gene candidates from the *C. tuberculosum* transcriptome, we aim to facilitate future research on this fascinating organism.

## Methods

### Data and code availability

All sequencing data generated from this study are available at European Nucleotide Archive (transcriptome data: accession PRJEB39919; genome data: accession PRJEB39919). Genome supported transcripts, transcriptome assemblies, annotations, and an example of metabolic pathway extraction are available on Github (https://github.com/martonelab/geneAnnotCalliarthronTranscriptome/).

### Experimental model and subject details

#### Specimen collection and sequencing

Two male, haploid specimens of *Calliarthron tuberculosum* were collected October 6, 2013, from Bluestone Point (48.81952, -125.1640), Bamfield, British Columbia, Canada and verified as haploid male specimens by microscopy. A portion of each collected sample was pressed and deposited into the UBC herbarium with voucher codes A89970 and A89985. Voucher codes can be queried at https://herbweb.botany.ubc.ca/herbarium/search.php?Database=algae for more information. Calcified intergenicula and non-calcified genicula from each individual were divided into two portions for data collection: either whole tissue (Sample I+G/PTM1 in the dataset) or calcified tissue only (Sample I/PTM2 in the dataset). Total RNA was extracted using the Spectrum Plant Total RNA kit (Cat # STRN50, Sigma-Aldrich) and sequenced on the Illumina HiSeq 2000 platform (paired-end 2x100bp, insert size ∼220bp).

#### Abbreviation of enzyme names

CAD, (hydroxy)cinnamyl alcohol dehydrogenase; SAD, sinapyl alcohol dehydrogenase; CCoAOMT, caffeoyl-CoA O-methyl transferase; CCR, (hydroxy)cinnamoyl-CoA reductase; C3’H, p-coumaroyl shikimate 3’-hydroxylase; C4H, cinnamate 4 hydroxylase; 4CL, 4-hydroxycinnamoyl-CoA ligase; COMT, caffeic acid O-methyltransferase; F5H, ferulic acid ⁄ coniferaldehyde ⁄ coniferyl alcohol 5-hydroxylase; HCT, hydroxycinnamoyl-CoA:shikimate hydroxycinnamoyl transferase; PAL, phenylalanine ammonia-lyase

#### Transcriptome assembly and annotation

Illumina sequence reads were assembled using Trinity with the *de novo* mode at default setting [72], independently for each anatomical sample (I+G/PTM1 ; I/PTM2 in the ENA database). A reference transcriptome was also assembled *de novo* using Trinity by independently combining the sequence reads generated from both samples. The assembled transcripts were annotated using Blast2GO [73]. Briefly, each transcript was searched against the NCBI RefSeq protein database (BLASTX, *E* ≤ 10^-5^), and its putative function was inferred based on the top protein hit and Gene Ontology (GO) terms. These proteins were then mapped onto the corresponding metabolic pathways in the Kyoto Encyclopaedia of Gene and Genomes (KEGG) database [74]. Identification of genes present in KEGG annotated pathways were extracted using the pathview package [75].

#### Filtering contaminant sequences in genome assembled data

To identify putative contaminant sequences in the genome assembly, each genome scaffold was searched (BLASTN) against a database of archaeal, bacterial and viral genome sequences retrieved from the NCBI RefSeq database. Sequences with a significant hit (E ≤ 10^-5^, covering > 50% of the query length) were considered putative contaminants and removed from the genome assembly. To identify broad differences in sequence characteristics, genomic scaffolds with and without transcriptomic support were compared for G+C content and transcript length (Fig S1B). Scaffolds with no transcript support and low recovery of eukaryotic genes (< 6% BUSCO or CEGMA recovery) were also identified as likely putative contaminants and removed from the genome assembly.

#### Genome annotation guided by transcriptome evidence

Repetitive elements in the genome assembly were identified and masked using RepeatMasker version open-4.0.6 [76]. To maximize recovery of transcript support for genome scaffolds, the transcriptomes (I+G/PTM1; I/PTM2 in the dataset) were mapped against the masked genome scaffolds using PASA v2.0.2 [77], and full-length coding sequences (CDSs) were predicted with TransDecoder v5.0.1 [72]. These CDSs represent the primary set of putative genes and were used as extrinsic hints to guide *ab initio* gene prediction using AUGUSTUS v3.2.1 [78] from the genome scaffolds.

#### HMM based gene candidate search

Monolignol biosynthesis gene candidates were identified from the *C. tuberculosum* transcriptomic dataset using Hidden Markov Model (HMM) based searches [79]. Transcriptomic sequence contigs were translated into all six reading frames using EMBOSS Transeq [80]. This amino acid database was used for subsequent sequence searches. HMM profiles used to search for homologs in the transcriptome were produced by aligning amino acid sequences of a given protein or protein family using MUSCLE [81] with no manual adjustment. The profiles were searched against the translated *C. tuberculosum* dataset in HMMER searches [79] to look for putative sequence homologs. Sequences more than 100 amino acids long were retained for subsequent analysis. These sequences were then searched against the *Arabidopsis* (GenBank taxid:3701) proteome using NCBI’s BLAST [82] to verify their closest homolog match (BLASTP, *E* ≤ 10^-30^).

#### Domain and motif comparison

The monolignol biosynthetic genes and their overall gene families contain sequence domains that influence protein shape and function. To compare these key domains, multiple sequence alignments (MSA) of candidate amino acid sequences from *C. tuberculosum* with their land plant counterpart protein were produced. Sequences were aligned using MUSCLE under default settings [81]. Key domains and motifs were chosen based on available literature and highlighted in the MSA as indicated in each figure legend. In each MSA, an asterisk (*) represents full conservation; and a period (.) represents sites with conservation >50%. Accession numbers can be found in Appendix S1.

#### Gene tree analysis

Gene trees were reconstructed for the candidate sequences of *C. tuberculosum* identified. For each gene tree analysis, sequence candidates from *C. tuberculosum*, the functionally demonstrated enzyme sequence from land plants, enzyme sequences from the overall protein family from land plants, and the top 20 sequences identified by NCBI BLAST using *C. tuberculosum* candidates as a query against the total database using default settings (BLASTP, *E* ≤ 10^-20^) were compiled. Land plant sequences identified to represent the functional gene and overall gene family were curated by a literature search. For each set of sequences, a multiple sequence alignment was performed using MUSCLE with default setting [81]. Sites with <80% coverage were removed using trimAl [83]. IQTree was used to search for the evolutionary model alignment under a BIC criterion [84, 85]. A maximum likelihood tree was reconstructed using IQTree [86], with node support calculated based on 1000 ultrafast bootstrap pseudoreplicates in IQTree [86]. A clade is considered strongly supported when bootstrap value ≥ 95%. FigTree was used to edit branch width and colors [87]. Accession numbers can be found in Appendix S1.

#### Generation of genome data as additional support for transcriptome data

Genome data of *C. tuberculosum* were generated using Illumina IIx platform (paired-end 2×150bp reads, insert size ∼350 bp). An overview of the summary statistics for the genome assembly can be found in Table S1. Adapter sequences were removed using Trimmomatic v0.33 [88] (LEADING:25 TRAILING:25 HEADCROP:10 SLIDINGWINDOW:4:20 MINLEN:50). The generated filtered sequence reads and the previously published genome data (GenBank accession #: SRP005182) generated using the 454 pyrosequencing platform [43] were used in a *de novo* genome assembly using SPAdes [89]. The 454 reads were treated as unpaired, single-end reads in the assembly process. This *de novo* assembly was further scaffolded with the transcriptome data using the L_RNA_Scaffolder [90]. Putative contaminant sequences were removed based on shared similarity against known genome sequences from bacterial, archaeal, and viral sources in NCBI RefSeq (BLASTN, *E* ≤ 10^-5^), and subsequently based on discrepancy in G+C content of the assembled scaffolds, and the recovery of core eukaryotic genes (CEGMA and BUSCO). Because the genome assembly is fragmented, genome scaffolds on which no transcripts were mapped were filtered out, yielding the final genome assembly (21,672 scaffolds, total bases 64.15 Mbp). These genome scaffolds were used as additional support for the transcriptome data. For the reference transcriptome (combined I+G/PTM1 ; I/PTM2), putative coding sequences were predicted based on alignment of the assembled transcripts against the genome scaffolds using PASA [77] and TransDecoder [72], from which the coded protein sequences were predicted.

#### Completeness of transcriptome and genome data

The completeness of the genome and transcriptome data was assessed by the recovery of core conserved eukaryote genes with the Core Eukaryotic Genes Mapping Approach (CEGMA) [91] and Benchmarking Universal Single-Copy Orthologs (BUSCO) [92] datasets. CEGMA and BUSCO datasets (eukaryote odb9 and Viridiplantae odb10) were independently used as query to search against the predicted proteins from the reference transcriptome (combined IG and IO) using BLASTP (*E* ≤ 10^-5^) and against the same transcriptome using TBLASTN (*E* ≤ 10^-5^). The core CEGMA and BUSCO proteins were also queried against the 21,672 genome scaffolds using TBLASTN (*E* ≤ 10^-5^).

## Key Resources Table

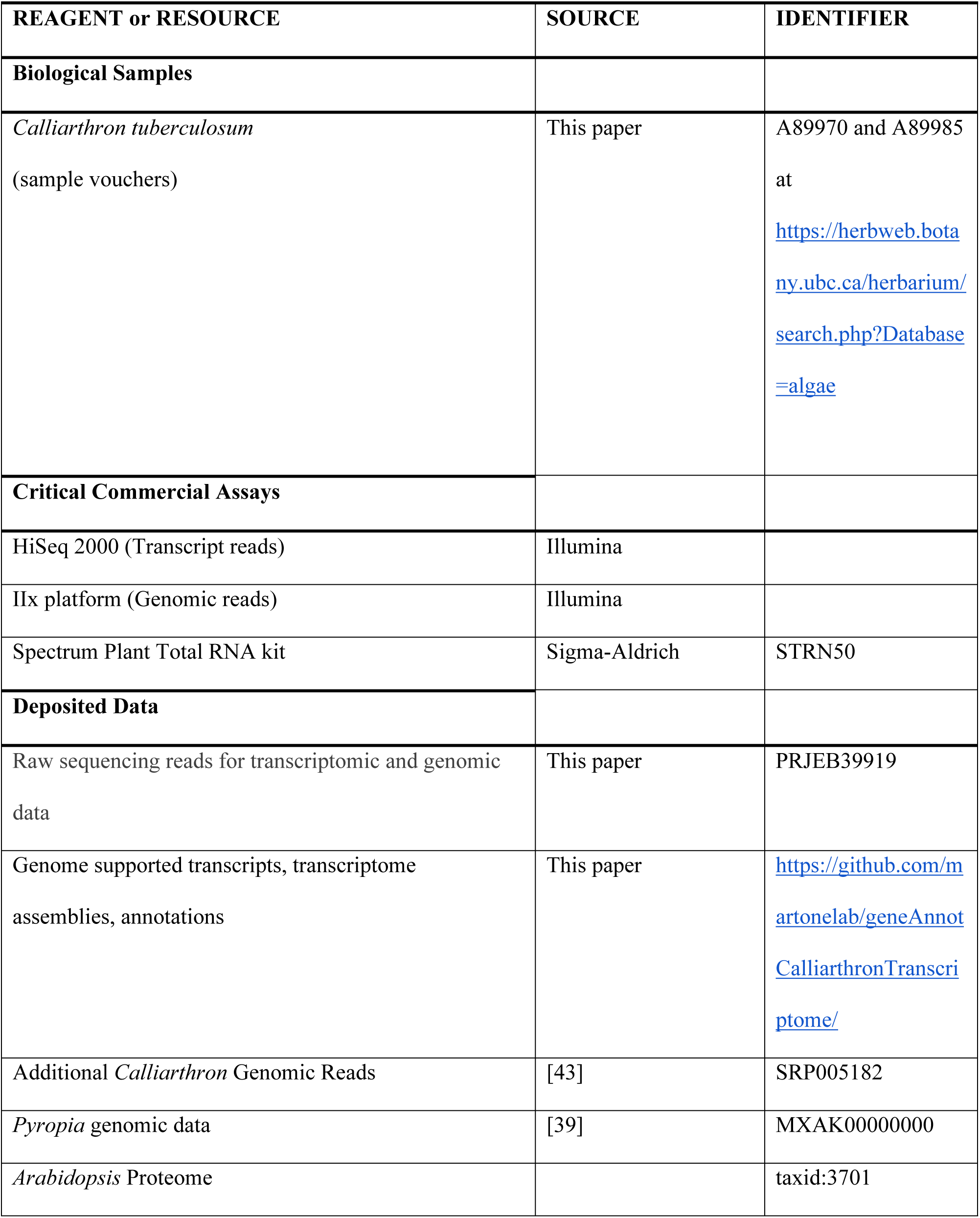
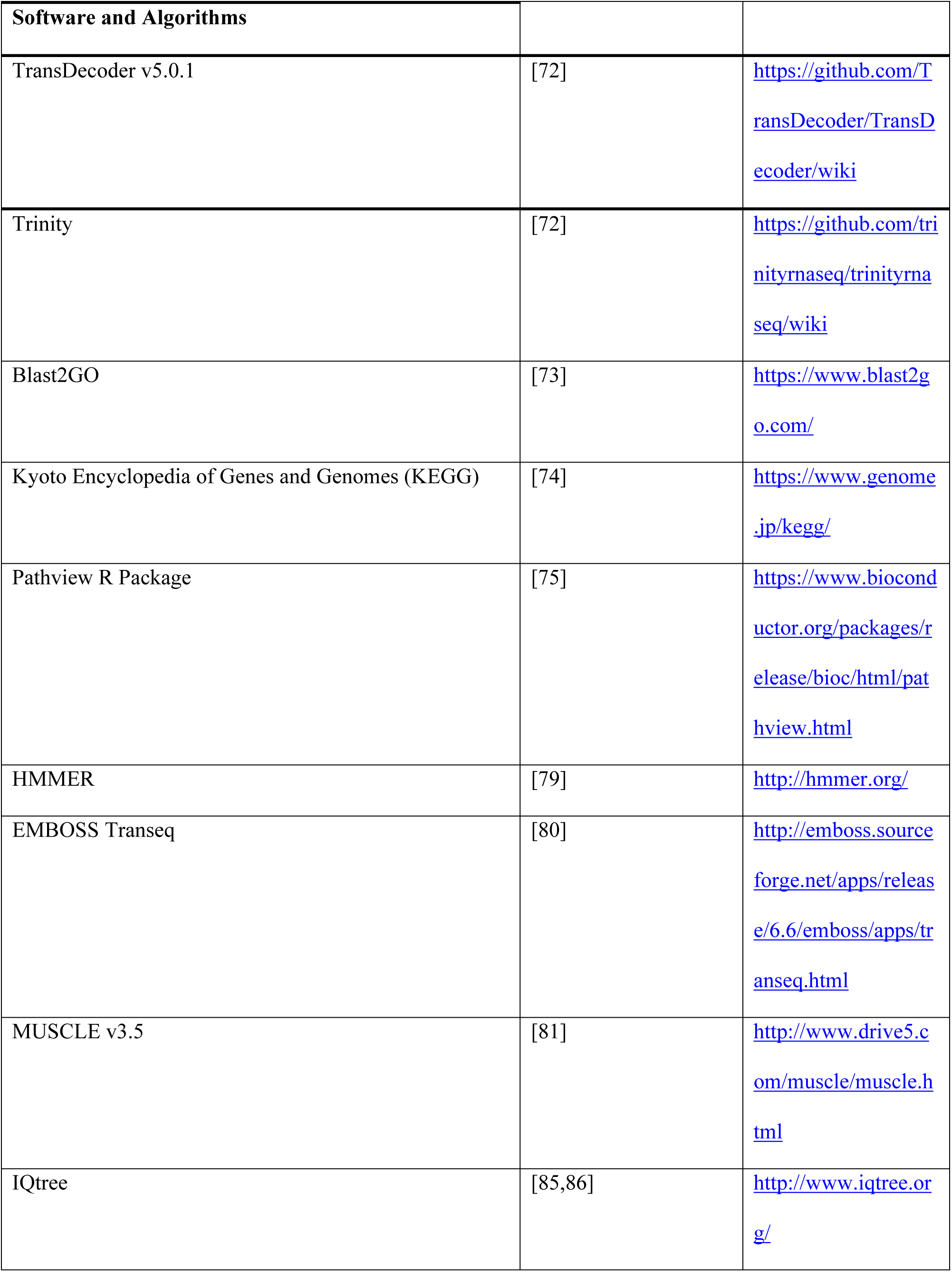
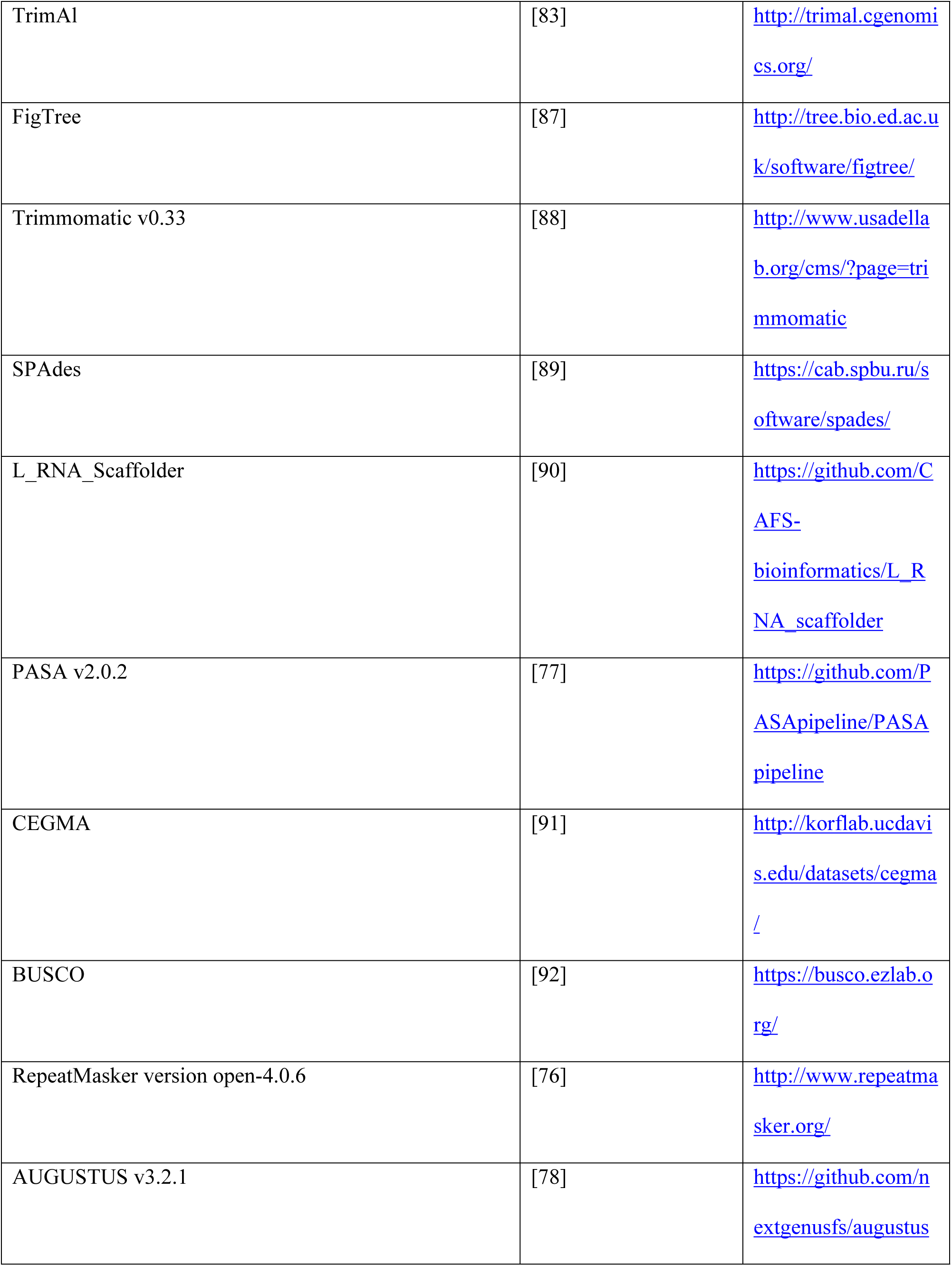

## Acknowledgements

We thank Dana Price and Debashish Bhattacharya (Rutgers University) for sequencing and preliminary analysis of the genome data, and Mike Thang (QFAB, Australia) for support in submitting sequence data to ENA. We thank the Osprey Ranch for supporting our writing retreats.

## Author Contributions

Conceptualization, J.X., K.H, and P.T.M; Methodology, J.X., K.H., M.A.L., C.X.C.; Investigation, A.M., J.X., K.H., M.A.L., C.X.C.; Visualization, J.X., E.J.; Writing - Original Draft, J.X. and P.T.M.; Review and Editing, J.X., K.H., M.A.L., E.J., C.X.C., P.T.M.; Funding Acquisition, P.T.M.; Supervision, P.T.M.

## Supporting Information

**Fig S1. Completeness of the *C. tuberculosum* transcriptome dataset.**

**(A)** Transcriptome sequences show high recovery of eukaryotic genes in CEGMA/BUSCO analysis. Percentage of genomic scaffolds with transcriptome support and transcriptomic scaffolds alone that share amino acid sequences with the core eukaryotic gene databases including CEGMA, BUSCO eukaryotic, and BUSCO Viridiplantae. Transcriptome encoded amino acid sequences were searched against the databases using BLASTP (orange) or TBLASTN (yellow), and genomic scaffolds were searched against the databases using TBLASTN (blue)

**(B)** Transcriptomic support of genomic data analyzed by GC content and transcript length. The distribution of GC content (above) against transcript lengths is shown for scaffolds with transcriptome support (blue) and scaffolds without transcriptome support (yellow) (right).

**Fig S2. C3H, C4H, F5H, P450 candidates from *C. tuberculosum* in relation to plants and other taxa.**

**(A)** Partial alignment of *C. tuberculosum* P450 candidates with C3H, C4H, and F5H from *A. thaliana*, and a novel F5H from *Selaginella moellendorffii*. Heme binding domain residues, secondary structure stabilizing K helix residues, PXRX, and the I-helix are indicated [8]. Sites with <80% coverage were removed. A strong candidate for beta-carotene synthesis is indicated with a triangle.

**(B)** Unrooted CYP450 maximum likelihood gene tree with *C. tuberculosum* (magenta dots) and additional taxa (Embryophyta – dark green, Chlorophyta – light green, Rhodophyta – red, Animalia and Opisthokonta – purple, Bacteria and Cyanobacteria – blue, Oomycota, Mycetozoa and Fungi – yellow, Ochrophyta – brown). Functionally demonstrated plant C3H, C4H, and F5H are labeled (+). Additional functional groups are labeled [9]. Ultrafastbootstrap values > 95 are marked by *. Model = VT+F+G4.

**Fig S3. CCoAOMT candidates from *C. tuberculosum* in relation to plants and other taxa.**

**(A)** Partial alignment of *C. tuberculosum* CCoAOMT sequence candidates with CCoAOMT from land plants. Substrate recognition residues (black triangle), divalent metal ion and cofactor binding residues (grey triangle), catalytic residues (back square), and the positively charged R220 necessary for substrate recognition (grey square) are indicated. Sites with < 70% coverage were removed.

**(B)** Unrooted maximum likelihood gene tree of biochemically characterized plant O-methyltransferases with *C. tuberculosum* (magenta dots) and additional taxa (Embryophyta – dark green, Chlorophyta – light green, Rhodophyta – red, Animalia and Opisthokonta – purple, Bacteria and Cyanobacteria – blue, Oomycota, Mycetozoa and Fungi – yellow, Ochrophyta – brown). Functionally demonstrated plant CCoAOMT are labeled (+). Additional functional groups are labeled [13]. Ultrafastbootstrap values > 95 are marked by *. Model = LG + G4. JMT, SAMT, and BAMT are closely related to OMTs.

**Fig S4. CSE candidates from *C. tuberculosum* in relation to plants and other taxa.**

**(A)** Partial alignment of *C. tuberculosum* CSE sequence candidates with CSE from land plants. Acyl transferase motifs (HX_4_D), lipase motifs (GXSXG) and active site residues (triangle) are indicated. Sites with < 70% coverage were removed.

**(B)** Unrooted maximum likelihood gene tree of *C. tuberculosum* CSE candidates (magenta dots) and additional taxa (Embryophyta – dark green, Chlorophyta – light green, Rhodophyta – red, Animalia and Opisthokonta – purple, Bacteria and Cyanobacteria – blue, Oomycota, Mycetozoa and Fungi – yellow, Ochrophyta – brown). Functionally demonstrated plant CSE are labeled (+). Additional functional groups are labeled. Ultrafastbootstrap values > 95 are marked by *. Model = VT+G4.

**Fig. S5. A visual representation of the *C. tuberculosum* sequences present in the starch and sucrose metabolism pathway from the KEGG based annotation.**

KEGG based annotation showing the starch and sucrose metabolic pathway with *C. tuberculosum* annotations highlighted. The gradient map in the top right corner indicates the level of transcription, with white and dark pink coloring representing absence and presence of expression respectively. The annotated map, number “00500”, was extracted in the provided R file using the pathview program.

**Table S1. Summary statistics for the *C. tuberculosum* genome assembly.**

Scaffolds are categorized as shared with either red algal (*Pyropia* yezoensis) genomic scaffolds, eukaryotic sequences, or other bacteria sequences based on sequence similarity.

**Table S2. Top hits against Arabidopsis thaliana (taxid:3702) using *Calliarthron* sequences as the search query (BLASTP).** Query sequence is indicated by contig number. Result hits are indicated by description (At tax ID 3702) and colored by overall alignment scores with red (>=200), pink (80-200), green (50-80), blue (40-50), and black (<40) that are most to least reliable scores in that order.

**Table S3. KEGG annotations of *Calliarthron tuberculosum* reads from the combined transcriptomic dataset.** Unique reads are represented by their contig identifier (contig_gene_isoform) and matched with their annotated KEGG based identifier (KO_identifier) and associated protein name.

**Table S4. Listed representation of the *C. tuberculosum* sequences present in starch and sucrose metabolism pathway from the KEGG based annotation.**

*C. tuberculosum* sequences were extracted from the KEGG based starch and sucrose metabolism pathway number “00500”. “KEGG Identifier” refers to the specific KEGG code for the gene, “Contig Name” refers to the sequence identifier from the *Calliarthron* transcriptome where the values represent the contig name_gene number_gene isoform and “Gene Name” refers to the gene acronym, the gene name, and its enzyme commission (EC) number. Sequences were extracted in the provided R file using the pathview program.

**Table S5. A list of calcification related gene candidates identified from KEGG-based annotations of the *C. tuberculosum* transcriptome.**

Calcification gene candidates were initially selected based on a literature search, and then *C. tuberculosum* sequences were identified manually from the KEGG based annotations (annotation file available on Github), thus this is not an exhaustive list. The genes are organized by their functional classification indicated as “overall function”, while “KEGG Identifier” refers to the specific KEGG code for the gene, “Contig Name” refers to the sequence identifier from the *Calliathron* transcriptome where the values represent the contig name_gene number_gene isoform and “Gene Name” refers to the gene acronym, the gene name, and its enzyme commission (EC) number.

